# Cross-modal perceptual enhancement of unisensory targets is uni-directional and does not affect temporal expectations

**DOI:** 10.1101/2021.03.06.434204

**Authors:** Felix Ball, Annika Nentwich, Toemme Noesselt

## Abstract

Previous studies demonstrated that redundant target stimuli can enhance performance due to multisensory interplay and interactively facilitate performance enhancements due to temporal expectations (TE; faster and accurate reactions to temporally expected targets). Here we tested whether other types of multisensory interactions – i.e. interactions evoked by temporally flanking irrelevant stimuli – can result in similar performance patterns and boost not only unisensory target perception (multi-vs. unisensory sequences) but also unisensory temporal expectations (expected vs. unexpected). To test our hypothesis, we presented sequences of 12 stimuli (10 Hz) which either consisted of auditory (A), visual (V) or alternating auditory-visual stimuli (e.g. A-V-A-V-…) with either auditory (AV(A)) or visual (AV(V)) targets. Participants had to discriminate target frequency which was unpredictable by temporal regularities (expected vs. unexpected target positions) and by stimulation sequence (A, V, AV(A), AV(V)). Moreover, we ran two experiments in which we presented redundant multisensory targets and manipulated the speed of the stimulation sequence (10 vs. 15 Hz stimulus trains) to control whether the results of Experiment 1 depended on sequence speed. Performance for unisensory targets was affected by temporally flanking distractors, with multisensory interactions selectively improving unisensory visual target perception. Yet, only redundant multisensory targets reliably affected TEs. Together, these results indicate that cross-modal facilitation of unisensory target perception in fast stimulus streams is uni-directional, but also differs from multisensory interactions evoked by redundant targets; more specifically, it appears to be context-dependent (task, design etc.) whether unisensory stimulation (unlike redundant target stimulation) allows for the generation of temporal expectations.

## 1. Introduction

Behavioural optimization is a crucial factor of our lives. For instance, to avoid accidents while driving, collision warning signals typically consist of redundant visual and auditory information to enable faster responses (Ho et al., 2007); when we talk with other people it is often easier to understand the other person when we process their lip movements in combination with their voice (Bernstein et al., 2004). These are just a few examples illustrating that multisensory interactions can facilitate behavioural responses and increase performance (more correct and faster responses; for review see Driver & Noesselt, 2008; Stevenson et al., 2014). A similar behavioural benefit is mediated by expectations; such expectations can be spatial (e.g. in right lane traffic one would expect a car approaching from the left side when crossing the street; for review on spatial attention see e.g. Carrasco, 2011) and/or temporal (e.g. it is easier to find a friend in a crowd when we know about their approximate time of arrival; for review on temporal attention see e.g. Nobre & Rohenkohl, 2014). Recently, it has been shown that redundant multisensory stimulation and temporal expectations (TEs) interactively enhance performance; as TE effects (higher accuracy and faster response times for targets at expected points in time) were enlarged in multisensory compared to unisensory conditions (Ball, Michels, et al., 2018; Ball, Fuehrmann, et al., 2018; Ball, Spuerck, et al., 2021). However, it is currently unknown whether only redundant target presentations (synchronously presented audio-visual targets) or also other forms of multisensory interplay (here the perceptual enhancement of targets by asynchronously flanked irrelevant stimuli) can cause such synergistic effects. Below, we will first introduce relevant studies on multisensory interplay before hypothesizing how these effects may interact with TEs.

To start with, a plethora of studies showed that an irrelevant stimulus can affect perception and reactions to a relevant target stimulus of a different modality (Lovelace et al., 2003; Maddox et al., 2015; Noesselt et al., 2010; Starke et al., 2020; Stein et al., 1996; Thorne & Debener, 2008; Van der Burg et al., 2008; Van Vleet & Robertson, 2006; Vidal et al., 2020; Vroomen & De Gelder, 2000). For instance, presenting an irrelevant auditory stimulus in combination with a visual target can result in increased perceived brightness of the visual target stimulus (Noesselt et al., 2008; Stein et al., 1996). Further, the time required to find a visual target stimulus among distractors is reduced and is hardly affected by the number of items in the search display when a spatially uninformative sound is concurrently presented (Van der Burg et al., 2008). Similarly, researchers showed that an alerting sound (un-as well as informative) can improve performance in visual conjunction search and reduce the amount of missing relevant targets in the attentional blink paradigm in patients with hemispatial neglect (Van Vleet & Robertson, 2006). Noesselt et al. (2010) suggested that effects of irrelevant sounds on visual perception are enlarged when stimulus intensity is lowered (i.e. stimuli close to the visual threshold), a finding which is in line with the theory of inverse effectiveness proposing that multisensory interactions are more likely to occur when stimulus perception is more difficult (Anastasio et al., 2000; Ball, Fuehrmann, et al., 2018; Colonius & Diederich, 2001, 2020; Ernst & Banks, 2002; Meredith & Stein, 1983, 1986; Stein & Meredith, 1993). While most previous studies focussed on the influence of the auditory on the visual modality there is also partial evidence that the visual modality can affect perception of relevant auditory targets (Lovelace et al., 2003; Starke et al., 2020; Thorne & Debener, 2008). For instance, Starke et al. (2020) showed that an uninformative and irrelevant visual stimulus improves performance in an auditory tasks (though see Vidal et al., 2020). Irrespective of the direction of cross-modal influences, all the effects mentioned above might be linked to activity changes of bimodal neurons which often exhibit supramodal superadditive activity (Meredith & Stein, 1986; Noesselt et al., 2010; Stevenson et al., 2007; Stevenson & James, 2009; Werner & Noppeney, 2010) but can also exhibit subadditive activity (Angelaki et al., 2009). While spatial proximity might be an important factor governing multisensory interaction under some circumstances (see e.g. Frassinetti et al., 2002), it may not be a necessary requirement for all types of multisensory interplay (for review and discussion see Spence, 2013), especially in temporal and non-spatial tasks (Ball, Michels, et al., 2018; Ball, Fuehrmann, et al., 2018; Ball, Spuerck, et al., 2021; Noesselt et al., 2005).

Of more importance for temporal tasks – as employed in this study – may be effects described by the ‘temporal rule’ (Stein & Meredith, 1993) and ‘Time-Window-of-Integration’ (TWIN; see Colonius & Diederich, 2004). Stein and Meredith (1993) suggested that multisensory interactions depend on close temporal proximity. Temporal integration is functionally important because light and sound propagate at different speeds and have different transduction times from their respective sensors to the cortex. In line with this idea, Frassinetti and colleagues (2002) showed that multisensory benefits do not only rely e.g. on spatial proximity but also on temporal proximity as multisensory enhancement was absent when stimuli were presented with a large inter-stimulus-gap (500 ms) as compared to being presented synchronously. Colonius and Diederich (2004) postulated that the time window works like a filter and that there is a limited time range in which two events of different modalities can be integrated or interact before they are processed and perceived as individual events. For audio-visual events, the interplay can e.g. depend on the duration, intensity and complexity of the stimuli as well as individual differences (see e.g. Boenke et al., 2009; Vatakis & Spence, 2006a, 2006b, 2007), with Stimulus-Onset-Asynchronies (SOA) of 0 to 200 ms still resulting in audio-visual cross-modal influences. A similar time window is found for visual-tactile events; according to Colonius and Diederich (2004), SOAs below 200 ms are optimal for the integration of visual-tactile stimulus events, resulting in faster reaction times for multisensory as compared to unisensory stimuli. In line with these reports, researchers reported TWINs of up to 200 ms, for instance, for the McGurk-effect (van Wassenhove et al., 2007). However, the TWIN can extend even further – up to 600 ms – depending on the specific study and thus, the utilized modalities, stimuli and tasks (Colonius & Diederich, 2011; Diederich & Colonius, 2015; Rach et al., 2011). Thus, these studies indicate that the width of the temporal interaction and integration window may not be generalizable. Nevertheless, there seems to be converging evidence that stimulus onset differences below 200 ms are beneficial for the potential interaction of stimuli of different modalities and that e.g. synchronous presentations often result in stimulus integration (single percept). In sum, the literature suggests that multisensory interactions and the related facilitation of performance are not restricted to synchronous presentations but can also occur when stimuli are presented asynchronously.

The research field of multisensory interactions is largely disconnected from that on TEs. Multisensory interactions are typically studied by manipulating modalities (uni-vs. multisensory events) while controlling for TEs by jittering targets’ onsets (Driver & Noesselt, 2008). TEs are typically studied by manipulating expectations about the temporal onset of events while keeping the modality constant (Nobre & Rohenkohl, 2014). TEs can be manipulated by embedding targets in rhythmic (predictable onset) versus arrhythmic (unpredictable onset) stimulus sequences or, for instance, by presenting targets with a higher likelihood after a short versus long cue-target interval (or vice versa). Note that TEs are not always assessed by an active judgement of time but rather by an orthogonal task. There, participants might use the temporal regularity to implicitly or explicitly shift attention to relevant moments in time (independent of the target modality) while the temporal regularity is non-predictive of the to-be-judged stimulus feature (e.g. judging target’s frequency; see Ball, Fuehrmann, et al., 2018; Ball, Michels, et al., 2018). Still, TEs have been shown to enhance response times as well as perceptual sensitivity for targets (Ball, Fuehrmann, et al., 2018; Ball, Michels, et al., 2018; Herbst & Obleser, 2019; Rohenkohl et al., 2012) although no information about the exact features of the upcoming stimulus was available. Such behavioural effects are in close resembles to effects of multisensory interplay.

Recently, we showed in a series of studies that the stimulation context (what is presented to participants) matters and that TE effects do not necessarily generalize across modalities and differ for uni-as compared to multisensory contexts (Ball, Fuehrmann, et al., 2018; Ball, Michels, et al., 2018). In our experiments, we presented sequences of 11 stimuli in which a single target was embedded on each trial. In each run, the target appeared with higher likelihood at an early position (3^rd^) or late position (9^th^). Participants discriminated the target frequency (lower or higher compared to distractors). While TEs were task irrelevant (the likely target onset provided no information about its frequency), they still facilitated performance (higher number of accurate and faster responses for expected trials). However, this effect was further interacting with the modality of the targets. Modality was manipulated by presenting auditory, visual or redundant audio-visual targets either in modality-specific sequences (pure auditory, visual or redundant audio-visual sequences) or in audio-visual sequences only (with unisensory auditory, unisensory visual or multisensory targets), thereby manipulating modality specific uncertainty. Independent of the uncertainty level, we observed that audio-visual target stimuli elicited robust TE effects while TE effect sizes in the unisensory conditions varied (Ball, Michels, et al., 2018). We further showed, that the TE effect was enlarged in the multisensory compared to the preferred, i.e. best (highest performance) unisensory condition (Ball, Fuehrmann, et al., 2018), providing a formal test of multisensory interactions (Stevenson et al., 2014) and indicating that multisensory interactions can affect TEs. Additionally, we were able to show that performance (independent of the TE effect) was elevated in the audio-visual compared to unisensory conditions (Ball, Fuehrmann, et al., 2018; Ball, Michels, et al., 2018). Thus, we also replicated well known effects from the literature showing superior performance for multisensory as compared to unisensory stimuli (Alais & Burr, 2004; Driver & Noesselt, 2008; Gondan et al., 2005; McDonald et al., 2000; Noesselt et al., 2007, 2010; Parise et al., 2012; Starke et al., 2020; Teder-Sälejärvi et al., 2005; Werner & Noppeney, 2010) but for the first time in a TE task. However, it is currently unknown whether the interaction of multisensory interplay and TEs can only be achieved by presenting redundant targets (synchronously presented audio-visual targets) or whether other forms of multisensory interplay (here the influence of an asynchronously presented irrelevant modality on the target modality) can also cause such synergistic effects.

Additionally, when unisensory targets were embedded in multisensory sequences we found that performance was diminished. Thus, instead of improving perceptibility of the unisensory target, a synchronously flanking irrelevant stimulus decreased performance while only the congruent stimulus (redundant multisensory target) improved performance. Note that this result might either be based on distractive effects (attention is drawn away from the relevant stimulus) or on attentional shifts: participants might have failed to monitor both stimulus streams and shifted attention from one modality to the other, thereby, potentially missing targets more often. The latter assumption would also be in line with the reduced TE effects in the unisensory condition as TE at least partially depends on perceiving the target at certain points in time. However, the detrimental effect of the flanking modality should be reduced when stimuli are presented asynchronously, i.e. they still may interact but are not fully integrated. In the present study, we therefore test whether unisensory target stimuli are more easily recognized in bimodal sequences when auditory (A) and visual (V) stimulus modalities are presented asynchronously (alternating for example A-V-A-V …). Further, by separating stimuli in time we test whether asynchronous presentation can result in multisensory facilitation and thus, potentially enlarges TE effects as compared to pure unisensory stimulations. Given our previous results (Ball, Fuehrmann, et al., 2018; Ball, Michels, et al., 2018) it is possible that TE for mere unisensory targets are always reduced to a minimum. However, if multisensory interplay is caused by temporally flanking irrelevant stimuli, we should find that performance in the multisensory condition is increased as compared to the unisensory trials. Further, this interplay might affect TEs, thereby increasing performance on expected trials in the multisensory condition.

## 2. Experiment 1

### 2.1. Methods

#### 2.1.1. Participants

We collected data from 36 participants. All participants provided written informed consent and declared to be free of neurological or psychiatric disorders and to have normal or corrected visual acuity. This study was approved by the local ethics committee of the Otto-von-Guericke-University, Magdeburg. Based on our previous studies (Ball, Fuehrmann, et al., 2018; Ball, Michels, et al., 2018), one participant was excluded due to a large response bias (using one response option in > 65 % of all trials) and another one due to low performance (performance in at least one condition < 25 % correct). We excluded another four participants due to technical problems with the equipment and stimulation, leaving 30 participants for analysis (mean age ± SD: 24.8 ± 4.8, # women: 17, # left-handed: 4).

#### 2.1.2. Apparatus

The experiments were programmed using the Psychophysics Toolbox (Brainard, 1997) and Matlab 2012b (Mathworks Inc.). Stimuli were presented on a LCD screen (22’’ 120 Hz, SAMSUNG 2233RZ) with optimal timing and luminance accuracy for vision researches (Wang & Nikolić, 2011). Resolution was set to 1650 x 1080 pixels and the refresh rate to 60 Hz. Participants were seated in front of the monitor at a distance of 102 cm (eyes to fixation point). Responses were collected with a wireless mouse (Logitech M325). Accurate timing of stimulation (<= 1 ms) was confirmed with a BioSemi Active-Two EEG amplifier system connected with a microphone and photodiode.

#### 2.1.3. Stimuli

Uni- or multisensory stimulus sequences (pure tones, circles filled with chequerboards, or a combination of both) were presented on each trial. Chequerboards subtended 3.07° visual angle and were presented above the fixation cross (centre to centre distance of 2.31°). Chequerboards were presented on a dark grey background (RGB: 25.5). The fixation cross (white) was presented 2.9° above the screen’s centre. Sounds were presented via headphones (Sennheiser HD 650). Note that we decided to use headphones as our previous research indicated that multisensory interplay effects in the present task might be enhanced when headphones are used (Ball, Fuehrmann, et al., 2018; Ball, Michels, et al., 2018).

Chequerboards and pure sounds were used as targets and distractors. The distractor frequencies were jittered randomly between 4.6, 4.9, and 5.2 cycles per degree for chequerboards and between 2975, 3000, and 3025 Hz for sounds. Visual and auditory target frequencies were individually adjusted to a 75 % accuracy level at the beginning of the experiment. Hence, targets – although the same type of stimulus (chequerboard/pure sound) – were either lower or higher in frequency compared to distractor frequencies. Furthermore, the intensities for both target and distractor chequerboards and sounds were varied randomly throughout the stimulus sequences. The non-white checkers were jittered between 63.75, 76.5, and 89.25 RGB (average grey value of 76.5 RGB). The sound intensities were jittered between 20 %, 25 %, and 30 % of the maximum sound intensity (average of 25 % = 52 dB[A]).

#### 2.1.4. Procedure

Participants were seated in a dark, sound-attenuated chamber. The experiment started with a short training (32 trials) to ensure that the participants understood the task and to get used to the stimulus material. Subsequently, a threshold measurement was used to adjust the target stimulus frequency of the unimodal modalities so that the participants answered correctly in 75 % of cases. Then the main experiment began (6 runs á 224 trials). For each trial, a sequence consisting of 12 stimuli was presented (see <u>Figure 1</u>for example sequences). Stimuli were presented for 50 ms followed by a gap of 50 ms. Note that we shortened the gap between stimuli as compared to our previous report (Ball, Fuehrmann, et al., 2018; Ball, Michels, et al., 2018) for two reasons: first, we increased the number of conditions (4 instead of 3) and early target positions (2 instead of 1) in the present experiment. Hence, smaller gaps and thus, faster presentations helped to keep the completion of the experiment below 3 hours. More importantly, previous research on temporal windows of multisensory interplay suggests that closer temporal proximity should increase the chance of multisensory facilitation. Thus, smaller temporal gaps should increase the potential influence of multisensory interactions.

**Figure 1:**
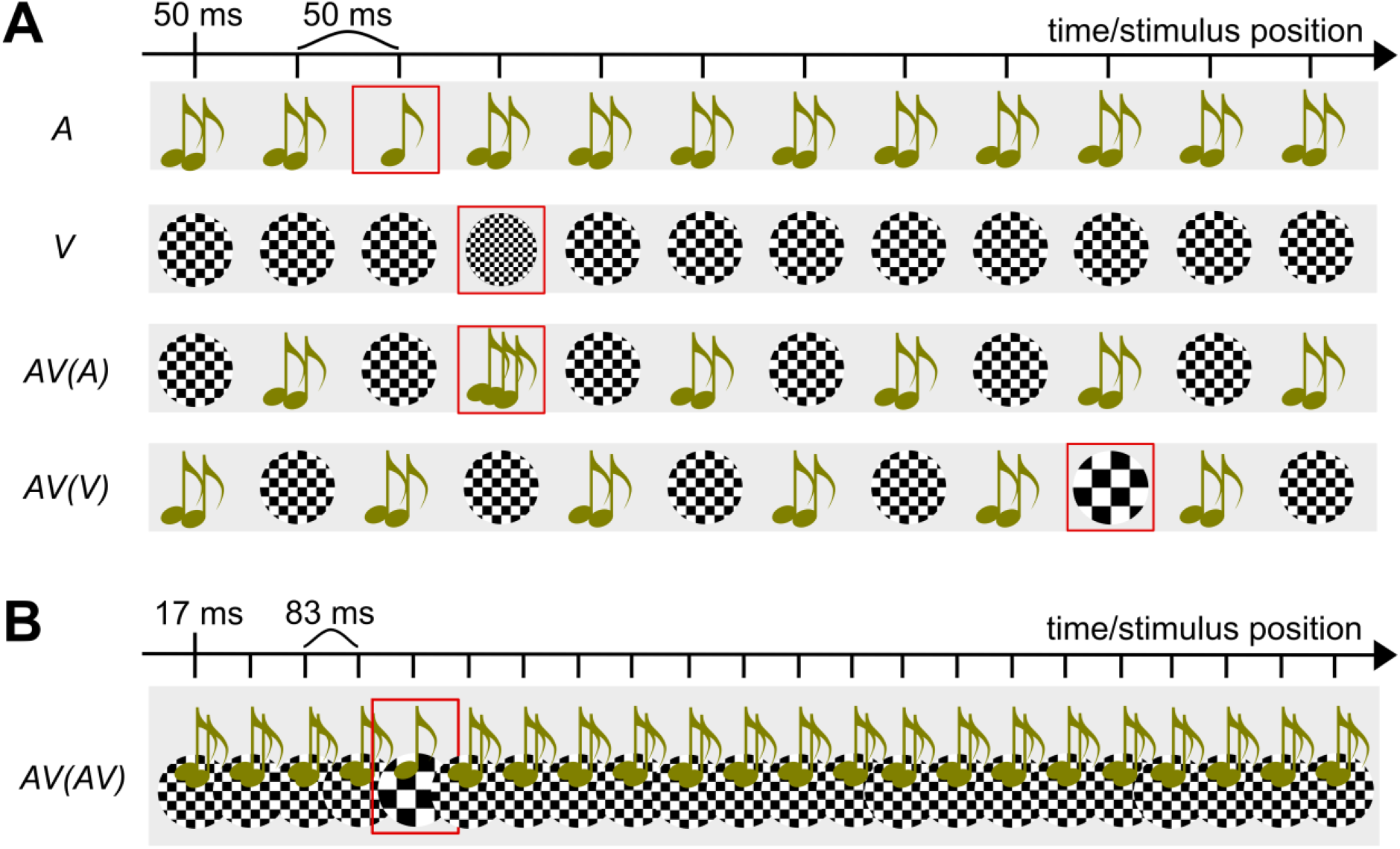
Schematic examples for stimulus sequences in Experiments 1 to 3. (**A**) Stimulus sequences used in Experiment 1. Here we presented always sequences of 12 stimuli (50 ms stimulus, 50 ms gap), with targets presented at the 3^rd^, 4^th^ or 10^th^ position (highlighted by red square which was not presented during the experiment). Targets had either a higher or lower frequency than distractors (auditory distractor frequency: 2975 - 3025 Hz, visual distractor frequency: 4.6 - 5.2 cycles per degree). Note that we only show examples and not all stimulus conditions (see main text for a description of all conditions). (**B**) Stimulus sequences in Experiment 2 were always multisensory and targets were redundant multisensory targets (both stimuli higher or lower in frequency as compared to distractors). Stimulus sequences contained 22 stimulus pairs (17 ms stimulus, 83 ms gap). Design of Experiment 3 is not shown directly but was like Experiment 2 with the difference that the gap between stimuli was reduced to 50 ms and we presented a total of 32 stimuli.

Sequences consisted either of just auditory (with auditory target), just visual (with visual target) or a mix of auditory and visual stimuli (with auditory [AV(A)] or visual target [AV(V)]). For the latter, modalities of stimulation were alternating and the sequence started either with an auditory or visual stimulus (A-V-A-V-… or V-A-V-A-…). Note that this procedure rendered the target stimulus unpredictable in multisensory sequences as it was not inferable from sequence onset. Thus, participants had to attend both modalities. Targets were either presented at positions 3 or 4 (early target) or 10 (late target) and had a higher or lower frequency than distractors. Each trial contained a target and participants were informed about this fact. Participants were asked to judge targets frequency (low or high) as correct and as fast as possible using a 2-alternative forced-choice procedure. Participants held the mouse with both hands, while placing their left/right thumbs on the left/right mouse buttons, respectively. Each button was used for one of the two response options (key bindings were counterbalanced across participants). The response recording started with the onset of the first stimulus of the sequence and ended 1500 ms after sequence’s offset. Only the first button press was recorded. The end of the response period was then followed by a 200 - 1200 ms inter-trial-interval. In case no button was pressed, the trial was repeated (mean of repeated trials across participants: 4.6 % ± 5.6 % SD).

Note that we kept expectations about target modality (A or V), sequence type (A, V, AV(A), AV(V)), early target position (3 or 4) and target’s frequency (low or high) unbiased by presenting them equally often in each run. However, we manipulated expectations about the general temporal position of targets (early or late within the sequence) by varying the likelihood of temporal positions across runs. In “expect early target” runs, early targets occurred in 86 % and late targets in 14 % of all trials. In “expect late target” runs, early targets occurred only in 43 % and late targets in 57 % of all trials. These two types of runs were presented alternatingly and the first run in each experiment was counterbalanced across participants. Note that these probabilities were based on our previous reports (Ball, Fuehrmann, et al., 2018; Ball, Michels, et al., 2018).

#### 2.1.5. Data analyses

For analyses, we used Matlab 2017b (Mathworks Inc.) and JASP (v 13.0.0). Note that accuracies were converted to d’ (measure of perceptual sensitivity; see Green & Swets, 1966) before analyses and mean response times were based on correct trials only. Further, in accord with our and other previous studies, we only analysed early target trials as late targets are always expected and may thus not require temporal attention (for a demonstration of the absence of late target TE effects see Ball, Fuehrmann, et al., 2018; Ball, Michels, et al., 2018; see also Jaramillo & Zador, 2011; Lange et al., 2003; Lange & Röder, 2006; Mühlberg et al., 2014 for similar a approach). The response window for trial inclusion was set to 150 – 2700 ms, with 2700 ms indicating the offset of the sequence plus the 1500 ms (response collection window) and RTs smaller than 150 ms indicating either responses prior to the target or insufficient target processing (Ball, Fuehrmann, et al., 2018; Ball, Michels, et al., 2018), resulting in a removal of 3 trials/participant on average. Further, performance was averaged across early target positions as these were balanced across conditions and to increase the trial number for estimation of performance in unexpected early trials.

Our first analysis focussed on general effects of TE and stimulation sequence type. To this end, we conducted two separate ANOVAs (one for d’ and one for mean RT) with factor TE (expected, unexpected) and sequence type (A, V, AV(A) and AV(V)). Based on our hypotheses, we would expect higher performance in the multisensory compared to the unisensory conditions as well as potentially enlarged TE effects (faster and more correct responses in expected trials) in the multisensory trials.

To test specifically for multisensory enhancement which cannot be inferred from the average performance values alone (see e.g. Ball, Fuehrmann, et al., 2018; Stevenson et al., 2014), we further identified the subject-specific “best” unisensory condition (highest performance and thus, preferred modality; see Ball, Fuehrmann, et al., 2018) and compared performance in this condition to its matching multisensory condition. Hence, if A was the best unisensory condition for the individual participant, performance was compared to the multisensory condition with auditory target; if V was the best unisensory condition we compared it to the multisensory condition with visual target. Note that we identified the preferred unisensory modality independently for d’ and mean RTs. Data were than analysed with two separate ANOVAs with within-subject factors TE (expected, unexpected) and modality (unisensory, multisensory) and between-subject factor modality preference (A, V).

If required, ANOVA results were Greenhouse-Geisser (p_GG_) corrected. Further, we used the false-discovery-rate correction (FDR; see Benjamini & Hochberg, 1995) to adjust results of all conducted post hoc tests (p_FDR_). Note that we used directional tests, to test for enhancement by multisensory stimulation. Only if these tests were non-significant, we used two-sided tests to test for possible declines in performance in the multisensory condition.

## 2.2. Results

### 2.2.1. Analyses of all conditions

Only the stimulation sequence type affected perceptual sensitivity (see Figure 2A; F(1.941,56.277) = 9.482, p_gg_ < .001, η_p_^2^ = .246). In particular (see Figure 2A), performance was enhanced in the multisensory condition with visual targets as compared to all other conditions (all ts > 2.814, all p_FDR_ = .019 or lower). Additionally, performance in both unisensory conditions exceeded performance in the multisensory condition with auditory targets (all ts > 2.474, all p_FDR_ = .032 or lower). The difference between the unisensory conditions was not significant (t(29) = .527, p = .709). Neither the TE effect nor its interaction with sequence type were significant (all F < 1.111, all p > .301).

**Figure 2:**
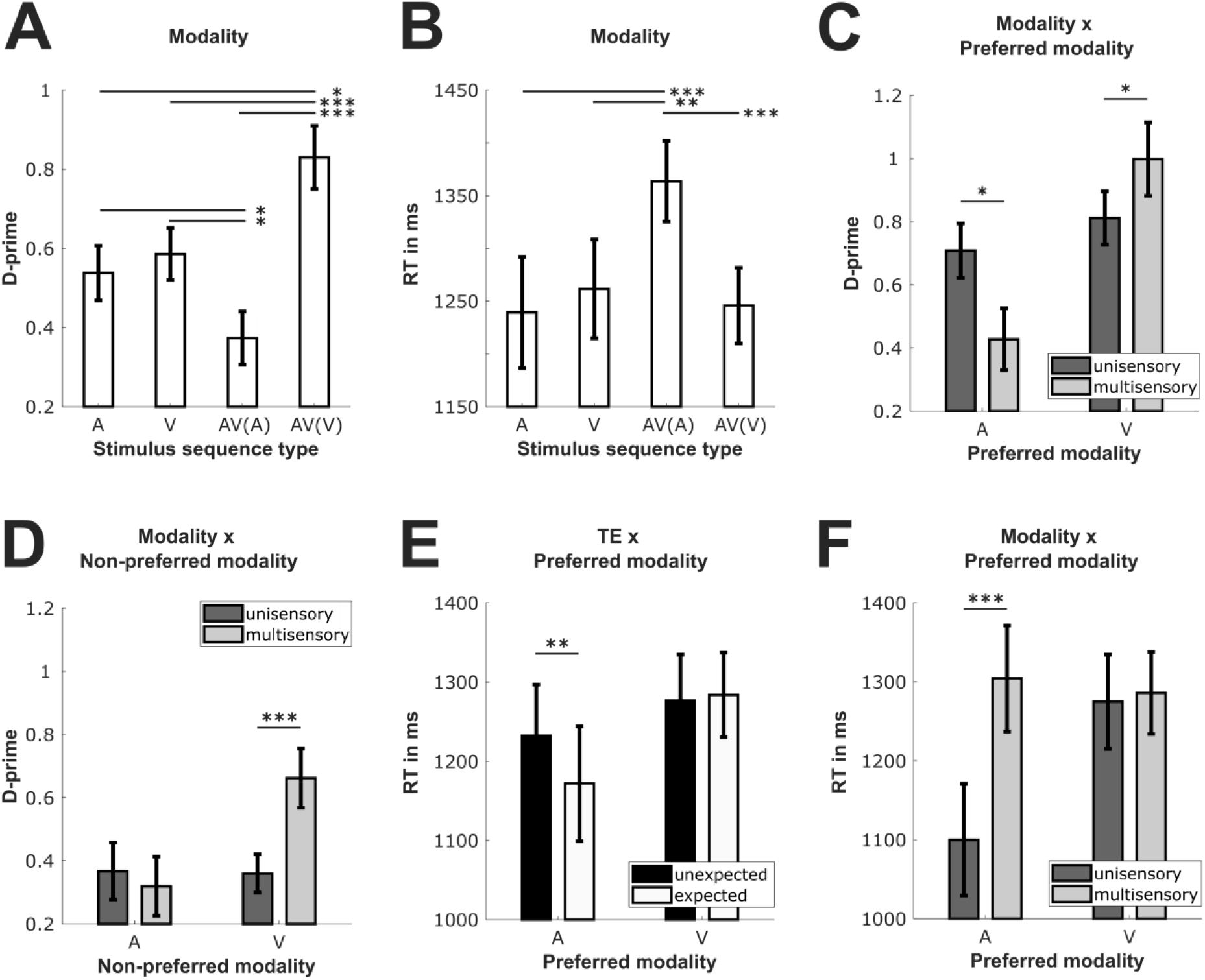
Performance effects in Experiment 1. Top row (from left to right): Main effect of modality for d’ (**A**) and mean RT (**B**; A – unisensory auditory, V – unisensory visual, AV(A) – multisensory with auditory target, AV(V) – multisensory with visual target), and interaction of modality and preferred modality for d’ (**C**; dark grey = unisensory trials, light grey = multisensory trials) **Bottom row (from left to right):** Interaction of modality and non-preferred modality for d’ (**D**; dark grey = unisensory trials, light grey = multisensory trials), interaction of TE and preferred modality for mean RT (**E**; black = unexpected trials, white = expected trials) and interaction of modality and preferred modality for mean RT (**F**; dark grey = unisensory trials, light grey = multisensory trials). Note that errorbars are standard errors and asterisks indicate significant differences between conditions (p < .001 [***], p < .01 [**], p < .05 [*]).

Response times were affected by TE, with responses being faster in the expected compared to the unexpected condition (mean_unexpected_ = 1294.4 ms, mean_expected_ = 1260.8 ms, F(1,29) = 6.611, p = .016, η_p_^2^ = .186). Further, we found again a modulation of performance by sequence type (see Figure 2B; F(2.263,65.641) = 8.373, p_gg_ < .001, η_p_^2^ = .224). Instead of an increase in performance, we only found reduced performance (i.e. increased response times) in the multisensory condition with auditory targets as compared to all other sequence types (all t > 3.774, all p_FDR_ < .002). All remaining post hoc tests were non-significant (all absolute t < .742, all p_FDR_ > .663) and the same was true for the interaction of TE and sequence type (F = 2.071, p = .11).

### 2.2.2. Test of multisensory enhancement

For perceptual sensitivity, 15 participants performed more accurately (i.e. preferred) in the unisensory auditory and 15 in the unisensory visual condition. For response times (RTs), 17 participants performed higher (i.e. preferred) in the unisensory auditory and 13 in the unisensory visual condition. For 16 participants, the preferred modality was identical between the two performance measures (9 participants with A and 7 with V preference).

The results for d’ showed a significant effect of preferred modality (F(1,28) = 7.977, p = .009, η_p_^2^ = .222) which further interacted with factor modality (see Figure 2C; F(1,28) = 11.804, p=.002, η_p_^2^ = .297). More importantly and in accord with our hypotheses, we found that for visual stimuli, performance was enhanced in multisensory trials (t(14) = 2.03, p_FDR_ = .048). However, for auditory trials, performance decreased in multisensory trials (t(14) = -2.8, p_FDR_ = .028). Note that this result does not imply multisensory inhibition in the auditory condition as this requires comparing the multisensory condition to cases when the auditory condition is the non-preferred (worst) unisensory condition. All remaining effects were non-significant (all F < .699, all p > .41).

For RT, the *TE* main effect approached significance (F(1,28) = 3.937, p = .057, η_p_^2^ = .123) and was further modulated by the factor *preferred modality* (see Figure 2E; F(1,28) = 6.203, p=.019, η_p_^2^ = .181): RTs in the expected condition were faster than in the unexpected condition but only if the preferred modality was auditory (t(16) = -3.279, p_FDR_ = .006) and not visual (t(12) = .354, p_FDR_ = .768). Further, we found a significant effect of modality (F(1,28) = 45.434, p < .001, η_p_^2^ = .619) which was – as for d’ – further modulated by factor *preferred modality* (see Figure 2F; F(1,28) = 36.425, p < .001, η_p_^2^ = .565). Importantly this interaction did not reveal any form of multisensory enhancement. If the visual modality was preferred, RTs did not significantly differ between uni- and multisensory trials (t(12) = -.502, p_FDR_ = .764). If the auditory modality was preferred, RTs were improved but in the unisensory and not multisensory condition (t(16) = -9.269, p_FDR_ < .001). All remaining effects were non-significant (all F < 2.502, all p > .125).

### 2.2.3. Test of multisensory inhibition

For both performance measures, we found no evidence of multisensory inhibition. Instead, the results suggested – at least for d’ – that perceptual sensitivity was enhanced by the multisensory stimulation even for the non-preferred modality. The results indicated a significant effect of *modality* (F(1,28) = 5.855, p = .022, η_p_^2^ = .173) which was further modulated by factor *non-preferred modality* (see Figure 2D; F(1,28) = 11.181, p = .002, η_p_^2^ = .285): here performance improved in the multisensory as compared to the unisensory condition – but again only for visual (t(14) = 4.435, t_FDR_ < .001) and not auditory stimuli (t(14) = -.608, p_FDR_ = .691). All remaining effects were non-significant (all F < 2.513, all p > .124).

Similar to the results for the preferred modality, we found that participants responded faster in expected compared to unexpected trials (meanunexpected = 1337.2 ms, mean_expected_ = 1301.2 ms, F(1,28) = 5.041, p = .033, η_p_^2^ = .153). Further, participants responded slower when the non-preferred modality was auditory (mean_A_ = 1233.3 ms, mean_V_ = 1431.6, F(1,28) = 6.344, p = .018, η_p_^2^ = .185). Crucially we found no evidence for multisensory inhibition as all remaining effects were non-significant (all F < 1.319, all p > .26).

### 3. Exp. 2 & 3: Effects of stimulation frequency on synchronous audio-visual targets

The results of Experiment 1 indicate the complete absence of TE effects for unisensory target stimuli even when cross-modal interplay was present and unisensory target perception was enhanced (i.e. when visual unisensory targets were temporally flanked by irrelevant sounds). Although the failure to induce robust TEs by unisensory stimulation has been observed before (Ball et al., 2020; Ball, Fuehrmann, et al., 2018; Ball, Michels, et al., 2018), there are several possibilities which could account for this outcome. Although performance was enhanced by cross-modal interplay for the visual target modality, performance (d’ ∼ 1) was slightly reduced as compared to performance in the redundant target condition in our previous report (d’ ∼ 1.2). However, note that the average performance in our previous studies did not determine the presence of TE effects which were also found for d’ < 1; thus, average performance differences cannot explain the absence of TE effects. Another possibility is that, in the present study, the likely time point of target occurrence was centred around 200 to 300 ms after sequence onset, instead of 400 ms (Experiments 1 to 4 in Ball, Michels, et al., 2018), thereby potentially decreasing TE effects. Such explanation is again unlikely supporting the pattern of results as we already showed (Experiment 6 in Ball, Michels, et al., 2018) that TEs can robustly alter performance even 200 ms after sequence onset and that this effect spreads across a larger time window (multiple target positions). Still, a last possibility is that the speed of the stimulus sequence itself affected the results. While we used a 5 Hz stimulus sequence in our previous studies, the present study utilized a 10 Hz stimulation sequence to facilitate potential multisensory interactions. This might have reduced TE effects as individual events were less separated and thus, targets might have been harder to identify. If this holds true, we should find similar results, namely, the absence of TE effects when using an audio-visual stimulation sequence with redundant audio-visual targets (both targets are higher or lower in frequency as compared to distractors) instead of unisensory targets.

To this end, we conducted two more experiments and analysed data of two independent groups of 30 participants each (Exp. 2: mean age ± SD: 24 ± 3.4, # women: 23, # left handed: 2; Exp. 3: mean age ± SD: 22.9 ± 3, # women: 19, # left handed: 1) after applying the same exclusion criteria as described for Experiment 1 (Exp. 2: one participants excluded, Exp. 3: three participants excluded).

To test the limits of creating TEs for the redundant audio-visual targets, we conducted two experiments in which we presented only sequences of synchronous audio-visual events (one target hidden among distractors). The main difference between experiments was that we varied the stimulation frequency to alter task difficulty (Exp. 2: 10 Hz; Exp. 3: 15 Hz). Task difficulty was additionally increased (as compared to Experiment 1) by presenting more stimuli within a sequence (Exp. 2: 22 stimuli with targets appearing at position 5 or 17; Exp. 3: 32 stimuli with targets appearing at position 7 or 25). As participants were naïve about the possible target positions, this increased the potential target positions two- and three-fold as (Exp. 1: 12 potential target positions). The increase in number of stimuli per sequence was achieved by shortening each stimulus presentation (distractors and target) to 17 ms, the minimal possible stimulus duration given our stimulation setup (the limiting factor was the monitor refresh rate of 60 Hz). Note that such short durations provide generally less time to process and identify target’s frequency as compared to Experiment 1. The two different stimulus sequence frequencies were achieved by using either an 83 ms (Exp. 2) or 50 ms (Exp. 3) inter-stimulus-gap. As in Experiment 1, we manipulated TE about the early and late target position across runs (6 runs á 56 trials) using the identical likelihoods of early and late targets. The remaining procedure was identical to Experiment 1. On average, three (Exp.3) to four (Exp. 2) trials/participant were excluded. Further, trial repetition was lower than in Experiment 1 (Exp. 2: 2.2 % ± 4.8 % SD; Exp. 3: 1.9 % ± 4.7 % SD). Again, we analysed differences between expected and unexpected early targets by conducting separate ANOVAs for each performance measure (d’ and RT) with repeated measures factor TE (unexpected, expected) and between-subject factor Experiment (Exp. 2, Exp. 3).

For both, d’ and RT, we only found a significant effect of TE. Perceptual sensitivity was increased in the expected compared to the unexpected condition (mean_unexpected_ = 1.213, mean_expected_ = 1.342, F(1,58) = 7.802, p = .007, η_p_^2^ = .119) and responses were also faster in the expected condition (mean_unexpected_ = 1305.1 ms, mean_expected_ = 1203.6 ms, F(1,58) = 37.147, p = < .001, ηp2 = .39). All remaining effects for both measures did not reach significance (all F < 2.163, all p > .147). Note that not only was performance, as measured by d’, exactly in the performance range of our previous studies (indicating differences between multisensory interplay affecting uni- and redundant multisensory targets); the results of Exp. 2 and 3 are also in line with the assumption that TE can be robustly created for audio-visual stimuli even when task difficulty is increased by various different factors and that the absence of TE effects is not driven by the stimulation sequence frequency but rather the unisensory target presentation itself.

## 4. Discussion

Previous studies demonstrated that redundant target stimuli can enhance performance due to synchronous multisensory stimulation and interacts with TE. Here we tested whether other types of multisensory interactions – i.e. interactions evoked by temporally flanking irrelevant stimuli – can result in similar performance patterns and boost not only TEs but also unisensory target perception. To test our hypothesis, we presented sequences of unisensory stimuli of the same modality (unisensory) or two alternating modalities (multisensory). We found multisensory enhancement to be present for unisensory targets but only when targets were presented visually (i.e. visual targets interacted with temporally flanking auditory distractors). Noteworthy, this type of enhancement was present irrespective of the subject-specific best unimodal condition. In contrast, when participants performed higher in the unisensory auditory condition, performance declined for auditory targets embedded in multisensory sequences. However, this finding was not indicative of multisensory inhibition as performance was equal between the unisensory and multisensory auditory target trials when participants performed lower in the auditory as compared to the unisensory visual condition. Moreover, although multisensory interactions enhanced visual target’s perceptibility (as indicated by performance changes), they did not significantly modulate TEs. TE effects were largely absent for unisensory stimuli and only present for the RT measure in the unisensory auditory condition. More importantly, although participants answered less accurate in the auditory conditions (lowest performance) they responded overall faster.

The results are mainly in line with our previous research showing that the learning of temporal regularities can be context-dependent and occurs with a higher likelihood for multisensory than unisensory targets (Ball et al., 2020; Ball, Andreca, et al., 2021; Ball, Fuehrmann, et al., 2018; Ball, Michels, et al., 2018; Ball, Spuerck, et al., 2021). Given the absence of TE effects in the unisensory conditions, we also replicated our findings that there seems to be an interactive effect of TEs and multisensory interactions, at least when induced by redundant targets. Here we additionally show that the multisensory TE effects (enhancement of response times and perceptual sensitivity) can occur for stimulation sequences with frequencies of up to 15 Hz. Note that in our previous studies (Ball, Fuehrmann, et al., 2018; Ball, Michels, et al., 2018; Ball, Spuerck, et al., 2021), we used a 5 Hz stimulus train leaving the possibility (although unlikely) that participants did not direct attention to a certain point in time but rather to a certain position in the stimulus sequence (i.e. they counted). However, the present results are based on such fast stimulation frequencies which render it impossible to count, thus, supporting our previous assumption that TE effects in the employed paradigm are based on shifts of attention in time and not position. Further, the results once more indicate that modality-specific TE effects for redundant multisensory targets are largely context-independent.

In the present study, we also found evidence for a different type of multisensory interplay, namely that unisensory performance can be enhanced by a temporally flanking irrelevant stimulus (see below for further discussion). However, this cross-modal enhancement of unisensory target perception in fast stimulation sequences appears to depend on the asynchronous presentation of irrelevant stimuli. Note that our previous studies showed that synchronously presented irrelevant stimuli decreased unisensory target perception (Ball, Fuehrmann, et al., 2018; Ball, Michels, et al., 2018; Ball, Spuerck, et al., 2021). Together these studies suggest that while synchronously presented irrelevant stimuli likely distracted participants, asynchronous presentation resulted – at least partially (visual only) – in cross-modal enhancement. Hence, asynchronous presentations may have reduced stimulus integration (single percept with contradicting information), potentially by keeping two separate stream representations (see e.g. Bregman & McAdams, 1994 for similar effects in auditory scene analysis), while interactions between streams still occurred. At the same time, however, this type of multisensory interplay appears to be insufficient to support reliable extraction of temporal regularities and thus, the build-up of TEs. This might be due to the fact that multisensory sequences did not only contain visual targets (enhanced by multisensory interplay) but also auditory targets (decreased performance) and that performance differences between both modalities reduced TE effects. However, the absence of TE effects in the unisensory modality might simply point to a limit of TE being invoked by unisensory stimuli. Please note that we do not propose that unisensory stimulations cannot induce TEs by learning of temporal regularities. There is in fact a rich literature that shows that unisensory temporal regularities can be learned and utilized (for review, see Nobre & Rohenkohl, 2014). However, these experiments are often restricted to analyses of reaction times, with accuracies being close to ceiling, indicating that the task at hand was rather easy. Noteworthy, we were able to show that such changes in reaction time do not necessarily indicate TEs (enhancement of response preparation or perceptual processing) but might simply imply changes in response strategies (Ball et al., 2020). Further, distractors are rarely used in unisensory TE experiments rendering the target easily detectable and even if distractors are used (e. g. when presenting a rhythmic sequence), the target is typically highlighted to increase detectability to a maximum (see e.g. Rohenkohl et al., 2012). All these factors show that typically, task difficulty is decreased to be able to investigate TEs in unisensory contexts. However, in real life, relevant information is not always highlighted or easily detectable. Hence, under more complex task regimes as the one employed (see also Ball, Fuehrmann, et al., 2018; Ball, Michels, et al., 2018), unisensory information might sometimes be insufficient to reliably extract information about the target and thus, temporal regularities. Thus, more context-dependent studies on TE are required, using different stimulus modalities, to investigate modality-specific effects and limits of the generation and utilization of TEs.

Note that in the remainder of the manuscript, we will focus on the d’ results as participants tended to focus on accuracy in the present study. This assumption is supported by response times being tightly linked to the end of the stimulus sequence and participants’ verbal reports after the experiment indicating that they typically waited till the sequence’s end to optimize their response (see also Ball, Michels, et al., 2018 for similar findings). Thus, differences in response times might be less readily interpretable in the current study. Further, response time effects in the analyses including factor ‘preferred modality’ rather point to speed-accuracy trade-offs than true response or perceptual facilitation. Overall, response times were faster when the auditory modality was the preferred modality (be it for the auditory modality in general or the expected trial condition). However, d’ was also lower in the preferred auditory as compared to the preferred visual modality. Hence, all ‘response time improvements’ were linked to generally lower accuracies in these conditions. This finding again shows that participants might utilize certain strategies to solve a task and that shifts in response times do not necessarily imply improved performance (see also Ball et al., 2020), unlike accuracy or d’ scores which always indicate performance shifts.

Turning to the bisensory results, i.e. higher perceptual sensitivity in the AV(V) sequence and lower perceptual sensitivity and slower response times in the AV(A) condition, these results can be linked to different phenomena reported in the multisensory literature. According to the modality appropriateness theory (Welch & Warren, 1980), performance and perception under multisensory stimulation is dominated by the sensory system that has a higher acuity and relevance for the task at hand. As a large portion of our daily experience is based on visual information, the visual modality is typically found to dominate other modalities (Koppen & Spence, 2007a, 2007b; Lukas et al., 2010; Sandhu & Dyson, 2012; Sinnett et al., 2008; Spence et al., 2011; Tomko & Proctor, 2017). Visual dominance is also found in spatial tasks (Ernst & Banks, 2002; Ernst & Bülthoff, 2004) while performance in temporal tasks is typically dominated by the auditory modality (Burr et al., 2009; Fendrich & Corballis, 2001; Noesselt et al., 2008; Recanzone, 2003; Repp & Penel, 2002). As we mainly employed a non-spatial task (time and stimulus frequency was relevant), the auditory modality might have affected the visual modality but not the other way around. It is for instance possible that the alternating presentation resulted in Sham’s illusion (Berger et al., 2003; Körding et al., 2007; Mishra et al., 2008; Rosenthal et al., 2009; Shams et al., 2000), thereby increasing the number of perceived visual stimuli induced by the flanking sounds. Thus, the target stimulus might have appeared e.g. twice. However, this would render the target stimulus less distinguishable from distractors as it would have been not unique anymore, thereby rather decreasing than increasing performance in the visual condition. Another possibility is that condition differences were mediated by ‘auditory driving’ (Gebhard & Mowbray, 1959; Shipley, 1964) or the ‘auditory ventriloquism’ effect (Aschersleben & Bertelson, 2003; Bertelson & Aschersleben, 2003; Ogulmus et al., 2018), both describing the effect that the visual modality is more likely to be temporally aligned with the auditory modality, either by biasing visual sequence frequency perception towards auditory’s sequence frequency or by ‘pulling’ visual stimuli in time towards the presentation of auditory stimuli. Hence, the auditory modality might have stabilized and enhanced the temporal perception of visual stimuli which might have enhanced performance (Driver & Noesselt, 2008). Yet, this stabilization, if present, did not affect TEs for visual targets. Further, if the auditory modality was dominant in multisensory trials, we should have rather found an increase for auditory than visual targets. Thus, all of the phenomena mentioned above lack to explain the sum of our results.

Also multisensory interactions can only partially explain our results. Here, successively presented stimuli in the multisensory sequences fell within the time window of multisensory integration (Diederich & Colonius, 2015). However, multisensory interactions are typically reported to enhance performance (Alais & Burr, 2004; Driver & Noesselt, 2008; Gondan et al., 2005; McDonald et al., 2000; Noesselt et al., 2007, 2010; Parise et al., 2012; Starke et al., 2020; Teder-Sälejärvi et al., 2005; Werner & Noppeney, 2010) which was only the case for the visual but not the auditory target in multisensory sequences. Thus, our results extend the literature of improvements of visual performance by irrelevant sounds (Maddox et al., 2015; Noesselt et al., 2010; Stein et al., 1996; Van der Burg et al., 2008; Van Vleet & Robertson, 2006; Vidal et al., 2020; Vroomen & De Gelder, 2000) by showing that such enhancement can also be achieved by asynchronously presented sounds embedded in fast stimulus sequences and when participants are oblivious about target’s modality (i.e. the multisensory sequence type was non-predictive of target’s modality). Noteworthy, the improvement in the visual condition was somehow enlarged when the visual condition was the non-preferred condition of the subjects. As ‘being the non-preferred condition’ likely implies that participants had more difficulties to perceive the visual target, our results are also in line with the principle of inverse effectiveness (Ball, Fuehrmann, et al., 2018; Meredith & Stein, 1983, 1986; Stein & Meredith, 1993). Yet, our results are not in agreement with the (albeit smaller) literature showing improvements of auditory performance by irrelevant visual stimuli (Lovelace et al., 2003; Starke et al., 2020; Thorne & Debener, 2008). The differences between the auditory and visual condition might be based on the stimulation context; in particular, the influence of auditory on visual activity in primary sensory areas appears to be a generalizable effect (Driver and Noesselt, 2008) – which would be supported by our results – while effects in the other direction are currently debated (for overview, see Starke et al., 2020). One necessary requirement for multisensory interactions induced by irrelevant visual stimuli might thus be the synchronous presentation of both stimuli as utilized in prior experiments (Lovelace et al., 2003; Starke et al., 2020; Thorne & Debener, 2008). However, this type of cross-modal influence would likely only be effective in single stimulus paradigms (see e.g. Starke et al., 2020) as we previously showed that for stimulation sequences, synchronously presented irrelevant stimuli reduce target perception (see e.g. Ball, Michels, et al., 2018).

Another explanation of our current results might be based on attention-related processes. For instance, participants might not have attended both stimulus modalities equally. Maybe the visual modality was attended more often, simply because participants preferred this modality or because it was more distracting. In turn, performance in the AV(A) condition would be reduced and performance in the AV(V) condition would be enhanced. As modality preferences (as measured by best unisensory performance) were balanced across modalities, the distraction by visual modality appears more likely and could be explained by the Colavita effect of visual dominance. Colavita (1974) reported that participants tend to report the visual and neglect the auditory stimulus when confronted with audio-visual presentations (for review see Spence et al., 2011). In his experiment, he presented random sequences of auditory and visual stimuli, with a few audio-visual events presented among them. On audio-visual trials, participants reported in 98 % of the cases the visual modality while being oblivious about the concurrent presentation of the auditory modality. The Colavita effect of visual dominance has been repeatedly addressed and investigated (Koppen & Spence, 2007a, 2007b; Lukas et al., 2010; Sandhu & Dyson, 2012; Spence et al., 2011; Tomko & Proctor, 2017). Further, Koppen and Spence (2007a) showed that the Colavita effect is stronger for stimuli in close spatial proximity and in an additional study (Koppen & Spence, 2007b), they showed that the effect occurs only for asynchronously presented stimuli with a temporal gap smaller than 150 ms, namely, before stimuli can be clearly temporally distinguished from one another. While the second requirement was fulfilled in our study (gaps of 50 ms), the first was not. Here, we presented sounds through headphones which likely prevented sounds from being spatially integrated and merged with the visual stimulus. Nonetheless, visual target perception was still enhanced which might indicate that the temporal factor is more important than the spatial factor for the emergence of the Colavita effect.

More importantly, neither the Colavita effect nor multisensory interactions alone can explain our findings. Rather, it is likely that a combination of both effects was at play. If the Colavita effect alone would underlie our results, we would expect that attention is directed prevalently to the visual condition. While such attentional biases can explain enhanced performance for visual as compared to auditory targets in the alternating sequences, they cannot explain the increased performance for visual targets in alternating vs. unisensory sequences. Even if one attends the visual condition in alternating sequence trials 100 % of the time, attention would be as strong as in the unisensory condition. Hence, we would expect similar performance scores between unisensory and alternating sequence trials. Further, attending the visual condition generally more often in alternating sequences should always result in performance decline for auditory targets in the alternating as compared to unisensory sequence trials (in the latter, auditory stimuli were attended 100 % of the time as no other modality was presented concurrently). However, the performance decline was only present when the unisensory auditory condition was the preferred modality (highest performance) and not the non-preferred modality. There were also no differences in performance between the unisensory auditory and visual condition and the preferred modality was balanced across participants. Thus, there was no intrinsic modality preference present in our current study. In sum, it is likely that our results were driven by an attentional bias in the alternating sequence condition that resulted in modality-specific enhancement due to multisensory interactions. Participants were more likely to attend the visual condition on multisensory trials and perception of the visual stimulus was enhanced by the flanking auditory stimulus. However, because participants likely attended the visual modality more often, it served potentially as some form of distractor when confronted with auditory targets, thereby hindering multisensory interactions and decreasing performance in the multisensory auditory condition. Note that a similar finding of concurrent multisensory enhancement for the visual modality and multisensory competition for the auditory modality was reported by Sinnett and colleagues. (2008) but only for response times and single stimuli (not stimulus streams). Here, we show that such cross-modal and asymmetrical multisensory interaction effects also manifest for perceptual sensitivity and in different stimulation contexts.

## 5. Conclusion

The results of this study demonstrate that temporal expectations are context-dependent and more reliably generated for multisensory target stimuli. Additionally, they indicate that presenting a distractor of a different modality asynchronously instead of synchronously within a stimulus stream, can facilitate unisensory target perception. However, even if unisensory target perception is enhanced by multisensory interactions (here the temporally flanking distractor), this increase in perceptibility appears to be insufficient for the extraction of temporal regularities. Thus, extraction of temporal regularities appears to depend on the stimulation context (isolated modality vs. mix of modalities across trials, single target vs. target embedded in distractors, target modality etc.) as well as whether and how different modalities compete for attention.

## Declarations

### Funding

This work was funded by the European Funds for regional Development (EFRE), ZS/2016/04/78113, Center for Behavioral Brain Sciences – CBBS.

## CRediT author statement

**Felix Ball:** Conceptualization, Methodology, Formal analysis, Investigation, Writing - Original Draft, Writing - Review & Editing, Visualization, Supervision, Project administration, Funding acquisition **Annika Nentwich:** Formal analysis, Writing - Original Draft, Writing - Review & Editing, Visualization **Toemme Noesselt:** Conceptualization, Methodology, Writing - Original Draft, Writing - Review & Editing, Supervision, Funding acquisition

## Ethics approval

This study was approved by the local ethics committee of the Otto-von-Guericke-University, Magdeburg.

## Consent to participate and publish

Informed consent was obtained from all individual participants included in the study.

## Competing interests

The authors declare no competing financial and non-financial interests, or other interests that might be perceived to influence the results and/or discussion reported in this paper.

## Open Practices Statement

The data and materials are available in the supplementary information files and upon request (e.g. codes). None of the experiments were preregistered.

## Data availability statement

The authors declare that the analysed data as well as potential supplementary analyses (not reported in the main manuscript) are available in the supplementary information files.

## Code availability

Data were analysed with JASP (freely available) and Matlab 2017b (Mathworks Inc.). Any relevant code is available upon request.

## Notes

### Competing Interest Statement

The authors have declared no competing interest.

### Summary of Updates

Added bioRxiv Links

## References

Alais, D., & Burr, D. (2004). The Ventriloquist Effect Results from Near-Optimal Bimodal Integration. Current Biology, 14(3), 257–262. https://doi.org/10.1016/j.cub.2004.01.029

Anastasio, T. J., Patton, P. E., & Belkacem-Boussaid, K. (2000). Using Bayes’ rule to model multisensory enhancement in the superior colliculus. Neural Computation, 12(5), 1165– 1187. https://doi.org/10.1162/089976600300015547

Angelaki, D. E., Gu, Y., & DeAngelis, G. C. (2009). Multisensory integration: psychophysics, neurophysiology, and computation. In Current Opinion in Neurobiology (Vol. 19, Issue 4, pp. 452–458). Curr Opin Neurobiol. https://doi.org/10.1016/j.conb.2009.06.008

Aschersleben, G., & Bertelson, P. (2003). Temporal ventriloquism: Crossmodal interaction on the time dimension: 2. Evidence from sensorimotor synchronization. International Journal of Psychophysiology, 50(1–2), 157–163. https://doi.org/10.1016/S0167-8760(03)00131-4

Ball, F., Andreca, J., & Noesselt, T. (2021). Context dependency of time-based event-related expectations for different modalities. Preprint at BioRxiv. https://www.biorxiv.org/content/10.1101/2021.03.06.434208v2

Ball, F., Fuehrmann, F., Stratil, F., & Noesselt, T. (2018). Phasic and sustained interactions of multisensory interplay and temporal expectation. Nature Scientific Reports, 8, 10208. https://doi.org/10.1038/s41598-018-28495-7

Ball, F., Groth, R.-M., Agostino, C. S., Porcu, E., & Noesselt, T. (2020). Explicitly vs. implicitly driven temporal expectations: No evidence for altered perceptual processing due to top-down modulations. Attention Perception & Psychophysics, 82, 1793–1807. https://doi.org/10.3758/s13414-019-01879-1

Ball, F., Michels, L. E., Thiele, C., & Noesselt, T. (2018). The role of multisensory interplay in enabling temporal expectations. Cognition, 170, 130–146. https://doi.org/10.1016/j.cognition.2017.09.015

Ball, F., Spuerck, I., & Noesselt, T. (2021). Minimal interplay between explicit knowledge, dynamics of learning and temporal expectations in different, complex uni- and multisensory contexts. Preprint at BioRxiv. https://www.biorxiv.org/content/10.1101/2021.03.06.434202v1

Benjamini, Y., & Hochberg, Y. (1995). Controlling the False Discovery Rate: A Practical and Powerful Approach to Multiple Testing. In Journal of the Royal Statistical Society. Series B (Methodological) (Vol. 57, pp. 289–300). WileyRoyal Statistical Society. https://doi.org/10.2307/2346101

Berger, T. D., Martelli, M., & Pelli, D. G. (2003). Flicker flutter: Is an illusory event as good as the real thing? Journal of Vision, 3(6), 406–412. https://doi.org/10.1167/3.6.1

Bernstein, L. E., Auer, E. T., & Takayanagi, S. (2004). Auditory speech detection in noise enhanced by lipreading. Speech Communication, 44(1-4 SPEC. ISS.), 5–18. https://doi.org/10.1016/j.specom.2004.10.011

Bertelson, P., & Aschersleben, G. (2003). Temporal ventriloquism: Crossmodal interaction on the time dimension: 1. Evidence from auditory-visual temporal order judgment. International Journal of Psychophysiology, 50(1–2), 147–155. https://doi.org/10.1016/S0167-8760(03)00130-2

Boenke, L. T., Deliano, M., & Ohl, F. W. (2009). Stimulus duration influences perceived simultaneity in audiovisual temporal-order judgment. Experimental Brain Research, 198(2–3), 233–244. https://doi.org/10.1007/s00221-009-1917-z

Brainard, D. H. (1997). The Psychophysics Toolbox. Spatial Vision, 10(4), 433–436. https://doi.org/10.1163/156856897X00357

Bregman, A. S., & McAdams, S. (1994). Auditory Scene Analysis: The Perceptual Organization of Sound. The Journal of the Acoustical Society of America, 95(2), 1177– 1178. https://doi.org/10.1121/1.408434

Burr, D., Banks, M. S., & Morrone, M. C. (2009). Auditory dominance over vision in the perception of interval duration. Experimental Brain Research, 198(1), 49–57. https://doi.org/10.1007/s00221-009-1933-z

Carrasco, M. (2011). Visual attention: The past 25 years. In Vision Research (Vol. 51, Issue 13, pp. 1484–1525). NIH Public Access. https://doi.org/10.1016/j.visres.2011.04.012

Colavita, F. B. (1974). Human sensory dominance. Perception & Psychophysics, 16(2), 409– 412. https://doi.org/10.3758/BF03203962

Colonius, H., & Diederich, A. (2001). A Maximum-Likelihood Approach to Modeling Multisensory Enhancement. In Advances in Neural Information Processing Systems (Vol. 14). www.uni-oldenburg.de/psychologie

Colonius, H., & Diederich, A. (2004). Multisensory interaction in saccadic reaction time: A time-window-of-integration model. Journal of Cognitive Neuroscience, 16(6), 1000– 1009. https://doi.org/10.1162/0898929041502733

Colonius, H., & Diederich, A. (2011). Computing an optimal time window of audiovisual integration in focused attention tasks: Illustrated by studies on effect of age and prior knowledge. Experimental Brain Research, 212(3), 327–337. https://doi.org/10.1007/s00221-011-2732-x

Colonius, H., & Diederich, A. (2020). Formal models and quantitative measures of multisensory integration: a selective overview. European Journal of Neuroscience, 51(5), 1161–1178. https://doi.org/10.1111/ejn.13813

Diederich, A., & Colonius, H. (2015). The time window of multisensory integration: Relating reaction times and judgments of temporal order. Psychological Review, 122(2), 232– 241. https://doi.org/10.1037/a0038696

Driver, J., & Noesselt, T. (2008). Multisensory Interplay Reveals Crossmodal Influences on “Sensory-Specific” Brain Regions, Neural Responses, and Judgments. In Neuron (Vol. 57, Issue 1, pp. 11–23). https://doi.org/10.1016/j.neuron.2007.12.013

Ernst, M. O., & Banks, M. S. (2002). Humans integrate visual and haptic information in a statistically optimal fashion. Nature, 415(6870), 429–433. https://doi.org/10.1038/415429a

Ernst, M. O., & Bülthoff, H. H. (2004). Merging the senses into a robust percept. In Trends in Cognitive Sciences (Vol. 8, Issue 4, pp. 162–169). Trends Cogn Sci. https://doi.org/10.1016/j.tics.2004.02.002

Fendrich, R., & Corballis, P. M. (2001). The temporal cross-capture of audition and vision. Perception & Psychophysics, 63(4), 719–725. https://doi.org/10.3758/BF03194432

Frassinetti, F., Bolognini, N., & Làdavas, E. (2002). Enhancement of visual perception by crossmodal visuo-auditory interaction. Experimental Brain Research, 147(3), 332–343. https://doi.org/10.1007/s00221-002-1262-y

Gebhard, J. W., & Mowbray, G. H. (1959). On discriminating the rate of visual flicker and auditory flutter. The American Journal of Psychology, 72, 521–529. https://doi.org/10.2307/1419493

Gondan, M., Niederhaus, B., Rösler, F., & Röder, B. (2005). Multisensory processing in the redundant-target effect: A behavioral and event-related potential study. Perception & Psychophysics, 67(4), 713–726. https://doi.org/10.3758/BF03193527

Green, D. M., & Swets, J. A. (1966). Signal Detection Theory and Psychophysics. Wiley.

Herbst, S. K., & Obleser, J. (2019). Implicit temporal predictability enhances pitch discrimination sensitivity and biases the phase of delta oscillations in auditory cortex. NeuroImage, 203, 116198. https://doi.org/10.1016/j.neuroimage.2019.116198

Ho, C., Reed, N., & Spence, C. (2007). Multisensory in-car warning signals for collision avoidance. Human Factors, 49(6), 1107–1114. https://doi.org/10.1518/001872007X249965

Jaramillo, S., & Zador, A. M. (2011). The auditory cortex mediates the perceptual effects of acoustic temporal expectation. Nature Neuroscience, 14(2), 246–251. https://doi.org/10.1038/nn.2688

Koppen, C., & Spence, C. (2007a). Spatial coincidence modulates the Colavita visual dominance effect. Neuroscience Letters, 417(2), 107–111. https://doi.org/10.1016/j.neulet.2006.10.069

Koppen, C., & Spence, C. (2007b). Audiovisual asynchrony modulates the Colavita visual dominance effect. Brain Research, 1186(1), 224–232. https://doi.org/10.1016/j.brainres.2007.09.076

Körding, K. P., Beierholm, U., Ma, W. J., Quartz, S., Tenenbaum, J. B., & Shams, L. (2007). Causal inference in multisensory perception. PloS One, 2(9), e943. https://doi.org/10.1371/journal.pone.0000943

Lange, K., & Röder, B. (2006). Orienting attention to points in time improves stimulus processing both within and across modalities. Journal of Cognitive Neuroscience, 18(5), 715–729. https://doi.org/10.1162/jocn.2006.18.5.715

Lange, K., Rösler, F., & Röder, B. (2003). Early processing stages are modulated when auditory stimuli are presented at an attended moment in time: an event-related potential study. Psychophysiology, 40(5), 806–817. https://doi.org/10.1111/1469-8986.00081

Lovelace, C. T., Stein, B. E., & Wallace, M. T. (2003). An irrelevant light enhances auditory detection in humans: A psychophysical analysis of multisensory integration in stimulus detection. Cognitive Brain Research, 17(2), 447–453. https://doi.org/10.1016/S0926-6410(03)00160-5

Lukas, S., Philipp, A. M., & Koch, I. (2010). Switching attention between modalities: Further evidence for visual dominance. Psychological Research, 74(3), 255–267. https://doi.org/10.1007/s00426-009-0246-y

Maddox, R. K., Atilgan, H., Bizley, J. K., & Lee, A. K. (2015). Auditory selective attention is enhanced by a task-irrelevant temporally coherent visual stimulus in human listeners. ELife, 2015(4), 1–11. https://doi.org/10.7554/eLife.04995.001

McDonald, J. J., Teder-Saälejärvi, W. A., & Hillyard, S. A. (2000). Involuntary orienting to sound improves visual perception. Nature, 407(6806), 906–908. https://doi.org/10.1038/35038085

Meredith, M. A., & Stein, B. E. (1983). Interactions among converging sensory inputs in the superior colliculus. Science (New York, N.Y.), 221(4608), 389–391. https://doi.org/10.1126/science.6867718

Meredith, M. A., & Stein, B. E. (1986). Visual, auditory, and somatosensory convergence on cells in superior colliculus results in multisensory integration. Journal of Neurophysiology, 56(3), 640–662. https://doi.org/10.1152/jn.1986.56.3.640

Mishra, J., Martinez, A., & Hillyard, S. A. (2008). Cortical processes underlying sound-induced flash fusion. Brain Research, 1242, 102–115. https://doi.org/10.1016/j.brainres.2008.05.023

Mühlberg, S., Oriolo, G., & Soto-Faraco, S. (2014). Cross-modal decoupling in temporal attention. The European Journal of Neuroscience, 39(12), 2089–2097. https://doi.org/10.1111/ejn.12563

Nobre, A. C., & Rohenkohl, G. (2014). Time for the Fourth Dimension in Attention. In A. C. Nobre & S. Kastner (Eds.), The Oxford Handbook of Attention (pp. 676–724). Oxford University Press.

Noesselt, T., Bergmann, D., Hake, M., Heinze, H. J., & Fendrich, R. (2008). Sound increases the saliency of visual events. Brain Research, 1220, 157–163. https://doi.org/10.1016/j.brainres.2007.12.060

Noesselt, T., Fendrich, R., Bonath, B., Tyll, S., & Heinze, H.-J. (2005). Closer in time when farther in space--spatial factors in audiovisual temporal integration. Brain Research. Cognitive Brain Research, 25(2), 443–458. https://doi.org/10.1016/j.cogbrainres.2005.07.005

Noesselt, T., Rieger, J. W., Schoenfeld, M. A., Kanowski, M., Hinrichs, H., Heinze, H.-J., & Driver, J. (2007). Audiovisual temporal correspondence modulates human multisensory superior temporal sulcus plus primary sensory cortices. The Journal of Neuroscience : The Official Journal of the Society for Neuroscience, 27(42), 11431–11441. https://doi.org/10.1523/JNEUROSCI.2252-07.2007

Noesselt, T., Tyll, S., Boehler, C. N., Budinger, E., Heinze, H.-J., & Driver, J. (2010). Sound-induced enhancement of low-intensity vision: multisensory influences on human sensory-specific cortices and thalamic bodies relate to perceptual enhancement of visual detection sensitivity. The Journal of Neuroscience : The Official Journal of the Society for Neuroscience, 30(41), 13609–13623. https://doi.org/10.1523/JNEUROSCI.4524-09.2010

Ogulmus, C., Karacaoglu, M., & Kafaligonul, H. (2018). Temporal ventriloquism along the path of apparent motion: speed perception under different spatial grouping principles. Experimental Brain Research, 236(3), 629–643. https://doi.org/10.1007/s00221-017-5159-1

Parise, C. V, Spence, C., & Ernst, M. O. (2012). When correlation implies causation in multisensory integration. Current Biology : CB, 22(1), 46–49. https://doi.org/10.1016/j.cub.2011.11.039

Rach, S., Diederich, A., & Colonius, H. (2011). On quantifying multisensory interaction effects in reaction time and detection rate. Psychological Research, 75(2), 77–94. https://doi.org/10.1007/s00426-010-0289-0

Recanzone, G. H. (2003). Auditory influences on visual temporal rate perception. Journal of Neurophysiology, 89(2), 1078–1093. https://doi.org/10.1152/jn.00706.2002

Repp, B. H., & Penel, A. (2002). Auditory dominance in temporal processing: new evidence from synchronization with simultaneous visual and auditory sequences. Journal of Experimental Psychology. Human Perception and Performance, 28(5), 1085–1099. https://doi.org/10.1037/0096-1523.28.5.1085

Rohenkohl, G., Cravo, A. M., Wyart, V., & Nobre, A. C. (2012). Temporal expectation improves the quality of sensory information. The Journal of Neuroscience : The Official Journal of the Society for Neuroscience, 32(24), 8424–8428. https://doi.org/10.1523/JNEUROSCI.0804-12.2012

Rosenthal, O., Shimojo, S., & Shams, L. (2009). Sound-induced flash illusion is resistant to feedback training. Brain Topography, 21(3–4), 185–192. https://doi.org/10.1007/s10548-009-0090-9

Sandhu, R., & Dyson, B. J. (2012). Re-evaluating visual and auditory dominance through modality switching costs and congruency analyses. Acta Psychologica, 140(2), 111– 118. https://doi.org/10.1016/j.actpsy.2012.04.003

Shams, L., Kamitani, Y., & Shimojo, S. (2000). Illusions: What you see is what you hear. Nature, 408(6814), 788. https://doi.org/10.1038/35048669

Shipley, T. (1964). AUDITORY FLUTTER-DRIVING OF VISUAL FLICKER. Science (New York, N.Y.), 145(3638), 1328–1330. https://doi.org/10.1126/science.145.3638.1328

Sinnett, S., Soto-Faraco, S., & Spence, C. (2008). The co-occurrence of multisensory competition and facilitation. Acta Psychologica, 128(1), 153–161. https://doi.org/10.1016/j.actpsy.2007.12.002

Spence, C. (2013). Just how important is spatial coincidence to multisensory integration? Evaluating the spatial rule. Annals of the New York Academy of Sciences, 1296(1), 31– 49. https://doi.org/10.1111/nyas.12121

Spence, C., Parise, C., & Chen, Y. C. (2011). The Colavita visual dominance effect. In The Neural Bases of Multisensory Processes (pp. 529–556). CRC Press. https://doi.org/10.1201/b11092-34

Starke, J., Ball, F., Heinze, H.-J., & Noesselt, T. (2020). The spatio-temporal profile of multisensory integration. European Journal of Neuroscience, 51(5), 1210–1223. https://doi.org/10.1111/ejn.13753

Stein, B. E., London, N., Wilkinson, L. K., & Price, D. D. (1996). Enhancement of perceived visual intensity by auditory stimuli: a psychophysical analysis. Journal of Cognitive Neuroscience, 8(6), 497–506. https://doi.org/10.1162/jocn.1996.8.6.497

Stein, B. E., & Meredith, M. A. (1993). The merging of the senses. The MIT Press.

Stevenson, R. A., Geoghegan, M. L., & James, T. W. (2007). Superadditive BOLD activation in superior temporal sulcus with threshold non-speech objects. Experimental Brain Research, 179(1), 85–95. https://doi.org/10.1007/s00221-006-0770-6

Stevenson, R. A., Ghose, D., Fister, J. K., Sarko, D. K., Altieri, N. A., Nidiffer, A. R., Kurela, L. R., Siemann, J. K., James, T. W., & Wallace, M. T. (2014). Identifying and quantifying multisensory integration: a tutorial review. Brain Topography, 27(6), 707–730. https://doi.org/10.1007/s10548-014-0365-7

Stevenson, R. A., & James, T. W. (2009). Audiovisual integration in human superior temporal sulcus: Inverse effectiveness and the neural processing of speech and object recognition. NeuroImage, 44(3), 1210–1223. https://doi.org/10.1016/j.neuroimage.2008.09.034

Teder-Sälejärvi, W. A., Russo, F. Di McDonald, J. J., & Hillyard, S. A. (2005). Effects of Spatial Congruity on Audio-Visual Multimodal Integration. Journal of Cognitive Neuroscience, 17(9), 1396–1409. https://doi.org/10.1162/0898929054985383

Thorne, J. D., & Debener, S. (2008). Irrelevant visual stimuli improve auditory task performance. NeuroReport, 19(5), 553–557. https://doi.org/10.1097/WNR.0b013e3282f8b1b6

Tomko, L., & Proctor, R. W. (2017). Crossmodal spatial congruence effects: visual dominance in conditions of increased and reduced selection difficulty. Psychological Research, 81(5), 1035–1050. https://doi.org/10.1007/s00426-016-0801-2

Van der Burg, E., Olivers, C. N. L., Bronkhorst, A. W., & Theeuwes, J. (2008). Pip and pop: Nonspatial auditory signals improve spatial visual search. Journal of Experimental Psychology: Human Perception and Performance, 34(5), 1053–1065. https://doi.org/10.1037/0096-1523.34.5.1053

Van Vleet, T. M., & Robertson, L. C. (2006). Cross-modal interactions in time and space: Auditory influence on visual attention in hemispatial neglect. Journal of Cognitive Neuroscience, 18(8), 1368–1379. https://doi.org/10.1162/jocn.2006.18.8.1368

van Wassenhove, V., Grant, K. W., & Poeppel, D. (2007). Temporal window of integration in auditory-visual speech perception. Neuropsychologia, 45(3), 598–607. https://doi.org/10.1016/j.neuropsychologia.2006.01.001

Vatakis, A., & Spence, C. (2006a). Audiovisual synchrony perception for music, speech, and object actions. Brain Research, 1111(1), 134–142. https://doi.org/10.1016/j.brainres.2006.05.078

Vatakis, A., & Spence, C. (2006b). Temporal order judgments for audiovisual targets embedded in unimodal and bimodal distractor streams. Neuroscience Letters, 408(1), 5–9. https://doi.org/10.1016/j.neulet.2006.06.017

Vatakis, A., & Spence, C. (2007). Crossmodal binding: Evaluating the “unity assumption” using audiovisual speech stimuli. Perception and Psychophysics, 69(5), 744–756. https://doi.org/10.3758/BF03193776

Vidal, M., Desantis, A., & Madelain, L. (2020). Irrelevant auditory and tactile signals, but not visual signals, interact with the target onset and modulate saccade latencies. PLOS ONE, 15(2), e0221192. https://doi.org/10.1371/journal.pone.0221192

Vroomen, J., & De Gelder, B. (2000). Sound enhances visual perception: Cross-modal effects of auditory organization on vision. Journal of Experimental Psychology: Human Perception and Performance, 26(5), 1583–1590. https://doi.org/10.1037/0096-1523.26.5.1583

Wang, P., & Nikolić, D. (2011). An LCD Monitor with Sufficiently Precise Timing for Research in Vision. Frontiers in Human Neuroscience, 5(85), 85. https://doi.org/10.3389/fnhum.2011.00085

Welch, R., & Warren, D. (1980). Immediate perceptual response to intersensory discrepancy. Psychological Bulletin, 88(3), 638–667. https://doi.org/10.1037/0033-2909.88.3.638

Werner, S., & Noppeney, U. (2010). Superadditive Responses in Superior Temporal Sulcus Predict Audiovisual Benefits in Object Categorization. Cerebral Cortex, 20(8), 1829– 1842. https://doi.org/10.1093/cercor/bhp248

